# Acute molecular and chronic vastus lateralis adaptations to lengthened partial versus full range of motion resistance training in previously trained males

**DOI:** 10.64898/2026.06.04.730150

**Authors:** Daniel L. Plotkin, Dakota R. Tiede, Tahran Gotla, Jaci Kelly, Mathis Rollin, Josh Queneua, Colin D. Wilborn, Marina Meyer-Vega, Valeria Robles-Cerdas, Adil Bashir, Ronald J. Beyers, Camila A. Esquivel, Christopher B. Mobley, Ryan Babl, Andreas N. Kavazis, Darren T. Beck, Harsimran S. Baweja, Christopher G. Vann, Paul A. Swinton, Lemuel W. Taylor, Michael D. Roberts

## Abstract

This study examined how lower-body lengthened partial (LP) versus full range of motion (FULL) resistance training affects acute post-exercise signaling, chronic hypertrophy, and cellular adaptations of the vastus lateralis (VL) muscle in resistance-trained men. Eight males (22±1 years old, 5.6±1.4 years training) completed a crossover study whereby VL biopsies were collected pre-exercise and 0, 3, and 24 hours following LP and FULL leg extension bouts for transcriptomic and anabolic signaling analyses (Experiment 1). Another 16 males (26±5 years old; 8.0±4.9 years training) completed an 8-week, twice-weekly lower-body intervention using a within-subject design (Experiment 2). One leg was assigned to FULL and the contralateral leg to LP training across three exercises (leg press, leg extension, and lying leg curl). Pre- and post-intervention outcomes included VL muscle cross-sectional area (mCSA) summed across five equidistant MRI-derived transverse slices and mid-thigh VL biopsy outcomes. As a secondary outcome, other hip and thigh muscles from Experiment 2 MRI scans were assessed. Condition × Time interactions for all outcomes were assessed using linear mixed-effects models. In Experiment 1, both conditions produced similar time-dependent changes in the VL transcriptome and anabolic (mTORC1 and Hippo) signaling, but minimal between-protocol interactions. In Experiment 2, VL summed mCSA significantly increased over time (mean change: 9.3 cm², 95% CI [6.8, 11.8], P<0.001), but there was no clear evidence of differential change between protocols (LP−FULL change: −1.4 cm², 95% CI [−6.1, 3.8], P=0.640). Additionally, no significant interactions existed for type I fiber CSA (P=0.476), type II fiber CSA (P=0.350), type I fiber myonuclei (P=0.813), type II fiber myonuclei (P=0.589), type I and II satellite cell number (P=0.102 and P=0.797, respectively), or total RNA content (P=0.537). Despite these null VL-centric findings, secondary Experiment 2 analyses provided some evidence that whole hamstring hypertrophy was greater following LP versus FULL (LP−FULL change: 3.9 cm², 95% CI [−0.2, 7.9], P=0.058). In conclusion, 8 weeks of LP and FULL resistance training broadly elicit similar acute and chronic VL responses in previously trained men, though secondary hamstring findings suggest that differential responses may depend on exercises included in the resistance training program.

## INTRODUCTION

Skeletal muscle hypertrophy is a primary adaptation to resistance training and is largely governed by the magnitude, distribution, and characteristics of the mechanical stimulus imposed on muscle [1]. Among the training variables that influence mechanical loading and subsequent hypertrophic outcomes, the muscle length and range of motion (ROM) through which an exercise is performed has received increasing attention. A growing body of evidence suggests that training at longer muscle lengths, either through positioning biarticular muscles in more lengthened joint configurations or by restricting repetitions to the lengthened portion of the movement (i.e., lengthened partial repetitions), may enhance hypertrophic outcomes relative to conventional full ROM training in specific muscles [2, 3].

Evidence for length-dependent hypertrophy in biarticular muscles exists for the hamstrings, which exhibit greater hypertrophy when trained in hip-flexion versus hip-neutral positions [4, 5]. Additional evidence exists for the rectus femoris, which exhibits greater hypertrophy when trained in a more hip-extended position [6], as well as the gastrocnemius when trained with the knee in the extended versus flexed position [7, 8]. The weight of the evidence also suggests that lengthened-biased ROM produces greater hypertrophy than shortened-biased ROM or full ROM training [2, 3]. Importantly, however, many of these full ROM comparators utilize exercises where the resistance challenge is predominantly shortened-biased, such as the calf raise and leg extension performed on conventional machines. While these findings are broadly consistent with a length-dependent hypertrophy advantage, several gaps in knowledge remain. These include whether findings generalize across muscles and resistance profiles, thresholds at which a greater lengthened bias may not confer greater hypertrophy, and whether such effects persist within full training programs incorporating exercises that emphasize varied muscle lengths.

Beyond these conceptual questions, several methodological gaps in the literature also limit interpretation. First, most studies have been conducted in untrained individuals. Second, most studies have used ultrasound-derived muscle thickness at one or two sites rather than MRI-based assessment of muscle cross-sectional area (mCSA) across the length of the muscle, which provides a more comprehensive characterization of the hypertrophic response. Third, no prior human work has combined whole-muscle MRI with biopsy-derived measures of fiber type-specific CSA (fCSA), satellite cell content, myonuclear number, and RNA content (a surrogate of ribosome content) when comparing ROM protocols, leaving the cellular mechanisms underlying any potential differential responses uncharacterized. Finally, the molecular mechanisms underlying any differential hypertrophic response, including acute transcriptomic and anabolic signaling responses to lengthened versus full ROM resistance exercise, remain uninvestigated.

Therefore, the purpose of this investigation was twofold. In our first experiment (termed Experiment 1 throughout), we sought to examine whether an acute bout of LP versus FULL unilateral leg extension exercise produced differential transcriptomic and anabolic signaling responses in the VL of resistance-trained men. In our second experiment (termed Experiment 2), we sought to examine whether 8 weeks of LP versus FULL lower-body resistance training produced differential VL hypertrophy and cellular adaptations in an independent cohort of resistance-trained men. To achieve our aims, we utilized various measures including MRI-derived mCSA across five equidistant slices and biopsy-derived measures of fCSA, satellite cell number, myonuclear number, and total RNA content, with secondary comparisons of non-VL hip and thigh mCSA. Given the knowledge gaps in this area of exercise physiology discussed above, we hypothesized that outcomes in both experiments would be similar between the LP and FULL protocols.

## METHODS

### Participants and Ethical Approval

All Experiment 1 and 2 procedures were conducted in accordance with the Declaration of Helsinki. The trials were not pre-registered given the hypertrophy-related outcomes of the project in younger healthy adults, and all participants provided written informed consent prior to enrollment and study procedures. Inclusion criteria for both experiments required: i) a minimum of 3 years of consistent resistance training experience with at least two lower-body sessions per week; ii) no lower-extremity musculoskeletal injury within the preceding 6 months; and iii) no contraindications to percutaneous skeletal muscle biopsies. Experiment 2 also contained the stipulation that no contraindications existed for MRI scanning. Experiment 1 occurred at the University of Mary Hardin-Baylor whereby 8 resistance-trained males between the ages of 18–35 years old were recruited from the local Belton, TX area (IRB protocol no. LT81725). Experiment 2 occurred at Auburn University whereby 18 resistance-trained males between the ages of 18–39 years were recruited from the local Auburn, AL area (IRB protocol no. STUDY00000282, Auburn University). Both experiments utilized convenience sampling without a priori sample size calculations. An overview of both experiments is presented in Figure 1.

**FIGURE 1.**
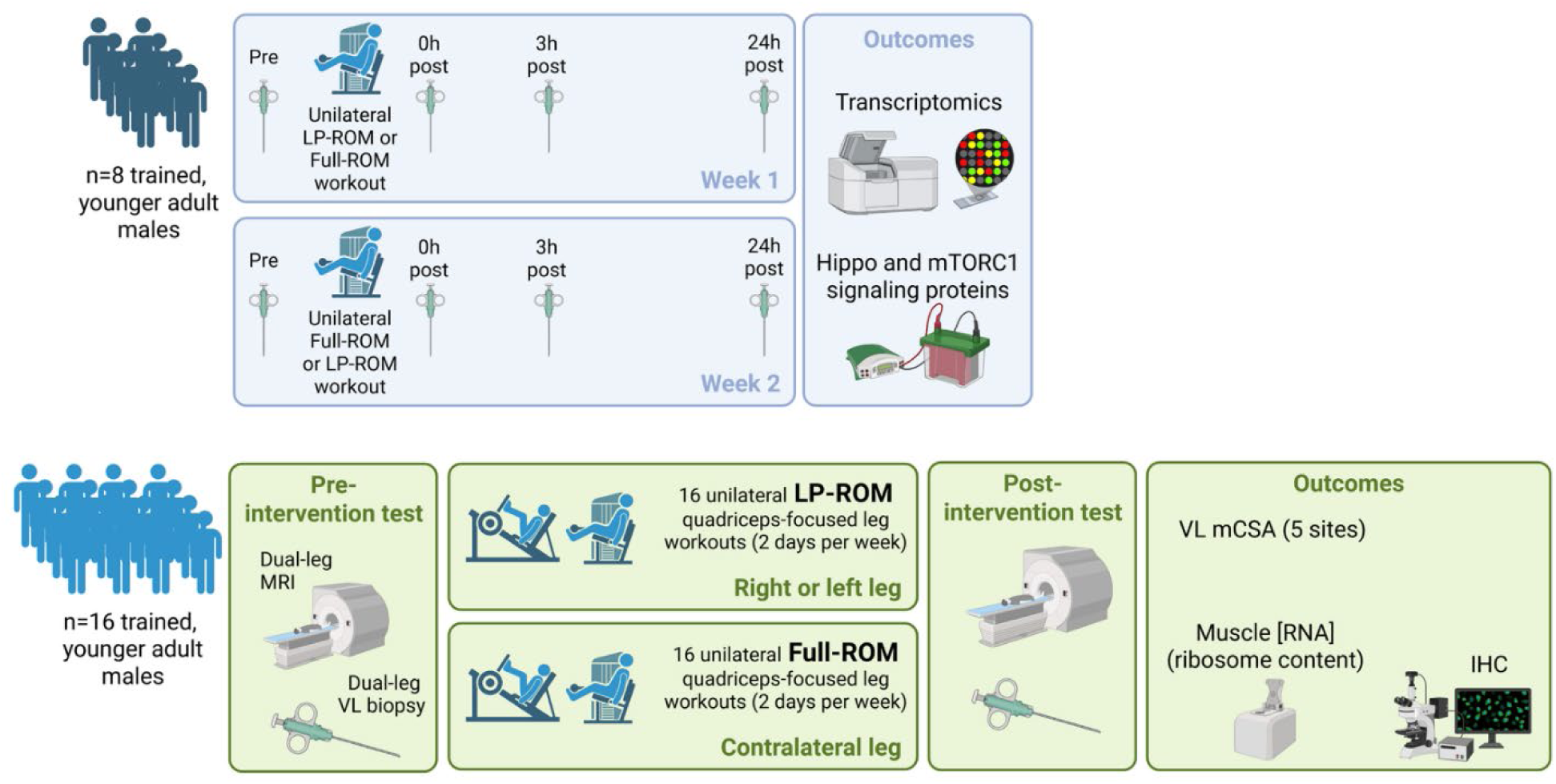
Study design overview. Legend: The upper panel depicts the Experiment 1 (acute) design, in which resistance-trained men completed a within-subject crossover with one leg performing LP-ROM and the contralateral leg performing Full-ROM unilateral leg extension across two separate weeks (counterbalanced order). VL biopsies were collected at PRE, 0h, 3h, and 24h post-exercise for transcriptomic and anabolic signaling analyses. The lower panel depicts the Experiment 2 (chronic) design, in which a separate cohort of resistance-trained men performed 16 unilateral LP-ROM (one leg) and Full-ROM (contralateral leg) quadriceps-focused workouts over 8 weeks (2 days/week), with pre- and post-intervention bilateral MRI and VL biopsy assessments. Abbreviations: Full-ROM, full range of motion; LP-ROM, lengthened partial range of motion; IHC, immunohistochemistry; mCSA, muscle cross-sectional area; VL, vastus lateralis.

### Experiment 1

Experiment 1 employed a within-subject crossover design in which participants completed two unilateral leg extension exercise bouts separated by one week. Participants first completed an informed consent and phenotyping visit including health history and a dual-energy x-ray absorptiometry (DXA) scan for body composition phenotyping purposes. A separate session established 10RM loads for each condition, conducted 3–5 days prior to the first training bout. Each leg was then randomly assigned to either FULL (100–0° knee flexion) or LP (100–50° knee flexion), and bouts were completed in counterbalanced order. Left and right limb assignment was block randomized using a computer random sequence generator to counterbalance for limb dominance, and exercise bouts were performed on a weighted stack leg extension machine (Model #20051, Cybex Eagle Nx Leg Extension, Medway, MA, USA). Participants arrived in an approximately 10-12-hour overnight fasted state between 0800 and 1200 for each visit. Each bout consisted of a standardized warm-up set consisting of 5 minutes on a stationary bike with low resistance followed by 5 unilateral leg extensor repetitions of ∼40% estimated one repetition maximum. Following a two-minute rest period, participants performed four unilateral leg extensor sets to concentric failure using an initial 10-repetition maximum (10RM) load, with adjustments made to maintain the 8–12 repetition range for subsequent sets, and 2 minutes of rest between sets. Participants were instructed to perform both the eccentric and concentric phases in a controlled fashion during earlier repetitions to minimize momentum, with the concentric phase encouraged to remain as explosive as possible as repetitions became more challenging. Vastus lateralis (VL) biopsies were obtained from the mid-thigh prior to (PRE) and immediately (0h), 3 hours (3h), and 24 hours (24h) after exercise as discussed in the next section. Participants were instructed to avoid lower-body exercise for at least 48 hours before each visit and to maintain similar dietary habits in the 48 hours prior to and the day of each bout.

### Muscle Biopsies

Mid-thigh VL biopsies were obtained using a 14-gauge needle with suction under aseptic conditions as previously described [9]. Three passes were collected yielding approximately 30 mg of tissue per biopsy. Tissue allocated was rapidly cleaned of blood, fat, and connective tissue, placed in cryogenic vials, flash frozen in liquid nitrogen within a 2-minute period following extraction, and stored at −80°C until shipment to Auburn University on dry ice for processing and analysis.

### Transcriptomics

RNA from VL tissue (∼15 mg) was isolated using Trizol (cat. no. 15596026; Thermo Fisher; Waltham, MA, USA) and associated reagents and centrifugation steps according to manufacturer’s instructions. RNA suspended in DEPC-treated water was shipped on dry ice to a commercial laboratory for RNA integrity checks and transcriptome-wide analysis using the Clariom S Assay Human mRNA array (North American Genomics; Decatur, GA, USA). Raw .CEL files were analyzed using Transcriptome Analysis Console (TAC) v4.0.2 (Thermo Fisher) with the *H. sapiens* genome for reference annotations. Two analyses were conducted including: i) pairwise comparisons to identify mRNAs altered from PRE within each bout whereby targets were considered significant at nominal P < 0.01 and log_2_-fold change ≥ |1.5| as we have previously performed with acute bout transcriptomic data [10], and ii) 2×2 repeated measures ANOVAs with the eBayes correction was used to identify mRNAs that differed between bouts over time, with interaction significance set at the same thresholds described above. Pathway analysis on resulting gene lists was performed using PANTHER v17.0 as previously described [10], whereby Gene Ontology (GO) Slim biological pathways were considered significant if pathway false discovery rate (FDR) values were less than 0.05. Pathways with less than 20 genes and/or not related to skeletal muscle physiology (e.g., disease-related pathways) were manually filtered out.

### Western Blotting

VL tissue (∼15 mg) was lysed on ice using general cell lysis buffer (Cell Signaling Technology; cat. no. 9803) with tight-fitting pestles. Total protein was quantified using a BCA protein assay kit (cat. no. A55864; Thermo Fisher) and spectrophotometer (Agilent Biotek Synergy H1, Thermo Fisher). Lysates were prepared for SDS-PAGE with 4× Laemmli buffer at equal protein concentrations (1 μg/μL) and 15 μL was loaded onto 4–15% Criterion TGX Stain-Free gels (Bio-Rad; Hercules, CA, USA). Electrophoresis was run at 200 V for 45–50 minutes. Proteins were transferred to PVDF membranes (Bio-Rad) for 2 hours at 200 mA, Ponceau stained and imaged (ChemiDoc Touch, Bio-Rad), and then blocked for 1 hour at room temperature with 5% nonfat milk powder in Tris-buffered saline with 0.1% Tween-20 (TBST, VWR International; Radnor, PA, USA), and incubated overnight at 4°C with primary antibodies (1:1000 in TBST with 5% bovine serum albumin) from a commercial vendor (Cell Signaling Technology; Danvers, MA, USA). These antibodies targeted phosphorylated (p-) LATS1 (Thr1079) (cat. no. 8654), p-YAP (Ser127) and p-TAZ (Ser89) (cat. no. 4911), p-mTOR (Ser2448) (cat. no. 5536), p-p70S6K (Thr389) (cat. no. 9234), and p-rpS6 (Ser235/236) (cat. no. 4858). The following day, membranes were washed for 15 minutes in TBST and incubated with a horseradish peroxidase-conjugated anti-rabbit secondary antibodies diluted 1:2000 in TBST with 5% BSA (cat. no. 7074; Cell Signaling Technology) at room temperature for 1 hour. Membrane development was performed using an enhanced chemiluminescent reagent (cat. no. ELLUF0100, Millipore Sigma; Burlington, MA, USA), and band densitometry was performed using a gel documentation system and associated analysis software (ChemiDoc Touch, Bio-Rad). Band densities for phosphorylated targets were normalized to corresponding Ponceau densities as we have previously performed with acute bout settings [11], and fold-change values were derived relative to the aggregate PRE mean of the FULL condition.

### Experiment 2

*Experimental Design.* Experiment 2 employed a within-subject contralateral limb design over 8 weeks using the 45-degree leg press, seated knee extension, and lying knee flexion exercises. One leg was assigned to FULL and the contralateral leg to LP for the duration of the intervention, and participants were block randomized by limb to counterbalance for limb dominance using blockrand package in R Studio [12].

Participants first completed an informed consent and phenotyping visit including health history screening, body composition assessment using bioelectrical impedance spectroscopy (SFB7, Impedimed; Carlsbad, CA, USA) and familiarization with the range of motion for each condition on the leg press. Three to five days thereafter, participants reported to the Auburn University MRI Research Center in a greater than 4-hour fasted state for pre-intervention lower-body MRI scanning. Participants were instructed to cease resistance training 72 hours prior to the initial MRI scan. One to two days following the MRI scan, pre-intervention bilateral VL biopsies were collected. Participants then completed 16 supervised resistance training sessions across 8 weeks (2 days/week). Eighty-seven to 96 hours after the final training session, participants returned for post-intervention MRI scanning, followed 1 to 2 days later by post-intervention bilateral VL biopsies. Participants were instructed to avoid lower-body resistance training outside of supervised sessions for the duration of the intervention.

### Resistance Training Protocol

Exercises were performed in the following order: unilateral leg press (Arsenal Strength Reloaded Linear Leg Press; Arsenal Strength, Lenoir City, TN, USA), unilateral leg extension, and lying leg curl (Life Fitness Signature Series; Life Fitness, Franklin Park, IL, USA). A 20-degree slant board (SquatWedgiez) was adhered to the footplate of the leg press using hook-and-loop tape to allow maximal knee flexion without losing heel contact. The seat back was positioned one setting from fully reclined for all participants. Both legs were trained to maximal knee flexion on the leg press, with all participants able to touch their calf to their hamstring at the bottom of each repetition. Foot placement was approximately hip width, positioned on the wedge to allow maximal hip and knee flexion to occur near simultaneously. The FULL leg did not lock out at the top of the movement to prevent offloading, and the LP leg extended to approximately the midpoint of this range. A metal device was held by a member of the research team at the LP endpoint to provide an audible cue when the sled plate reached its target position. Unilateral leg extension was performed through either FULL (100–0° knee flexion) or LP (100–50° knee flexion) ranges of motion, monitored visually by a researcher, with seat back positioned for 100° of hip flexion. Lying leg curl was performed through the full individualized ROM for the FULL leg and 5–10° beyond half of the individualized ROM for the LP leg to account for greater gastrocnemius involvement in the early portion of the range; the LP endpoint was enforced via a physical block held by a staff member. All ROM endpoints were visually assessed by trained research staff rather than mechanically measured, a pragmatic decision intended to reflect real-world training conditions. Participants were instructed to perform both the eccentric and concentric phases in a controlled fashion during earlier repetitions to minimize momentum, with the concentric phase encouraged to remain as explosive as possible as repetitions became more challenging. Leg press was performed to volitional failure, defined as the point at which the participant believed the next repetition could not be completed or would require compromised form. The leg extension and leg curl exercises were performed to concentric failure, defined as the inability to complete the assigned range of motion. Sets and rest periods were exercise-specific, and legs were alternated between sets in the following fashion: i) leg press progressed from 3 sets (sessions 1–9) to 4 sets (sessions 10-16) with 1.5 minutes of rest between sets, ii) leg extension progressed from 3 sets (sessions 1–3) to 4 sets (sessions 4 onward) with 1 minute of rest between sets, and iii) leg curl progressed from 5 sets (sessions 1–5) to 6 sets (sessions 6-16) with 1 minute of rest between sets. For each set, 8-12 reps were completed and the load was adjusted to reach the target intensity (volitional or concentric failure), within the target rep range. As such, loads were progressed throughout the intervention. The leg that was trained first in each exercise was alternated each session.

### Leg press kinematics

ROM adherence was confirmed during session 10 of 16, in which participants were fitted with inertial measurement units (IMUs; Noraxon Ultium Motion, Noraxon U.S.A., Scottsdale, AZ, USA) on each shin, each thigh, and the lumbosacral junction to verify that each leg was trained within its assigned range of motion via wireless transmission to myoRESEARCH software (Noraxon U.S.A). Raw joint angle data exported from myoRESEARCH were processed in R (version 4.4.1; R Core Team, 2024). Knee and hip flexion signals were each smoothed using a 7-point centered moving average filter prior to repetition detection. Individual repetitions were identified from peaks in the smoothed knee signal using a minimum peak separation of 0.20 seconds. The first and last repetitions of each set were excluded when four or more repetitions were detected to minimize the influence of transition periods. All processed files were visually inspected to confirm accurate repetition detection. Peak flexion, minimum flexion, and ROM were extracted from each retained repetition and averaged across repetitions within each set, then across sets within each leg.

### Magnetic Resonance Imaging

MRI scanning was performed on a 3T SkyraFit system (Siemens, Erlangen, Germany) at the Auburn University MRI Research Center. Participants were positioned prone on the patient table with an approximately 5-minute latency period before scanning. A T1-weighted turbo spin echo pulse sequence was used (repetition time: 1,400 ms; echo time: 23 ms; in-plane resolution: 0.9 × 0.9 mm²) to obtain transverse image sets spanning the top of the ilium to the patella with a 2-mm slice thickness and no inter-slice gap. All imaging was performed blinded to training conditions by the same investigator (R.J.B.). DICOM files were preprocessed in OsiriX MD (Pixmeo, Geneva, Switzerland) and imported into 3D Slicer (Brigham and Women’s Hospital, Boston, MA, USA), where the polygon tool was used to manually trace VL borders (primary study outcome) and other leg and hip muscle borders (secondary study outcome) on each image. Thigh image quantification was standardized as follows: i) the most proximal slice was defined as the first axial image in which the gluteal musculature was no longer visible, ii) the most distal slice was defined as 10 slices (2 cm) proximal to the final image in which the RF was no longer visible, and iii) three equidistant slices between these points were quantified, yielding five equidistant slices at both PRE and POST. Total VL and other muscle mCSA values for thigh musculature were computed as the sum of mCSA values across slices. Gluteal mCSA was assessed spanning 15 slices from the most proximal to most distal extent of the visible gluteal musculature, with 3 equidistant slices between these endpoints yielding five equidistant slices per timepoint. One participant was excluded from gluteus maximus analyses as the proximal muscle boundary was not visible within the field of view. All participants were missing the most proximal biceps femoris short head (BFsh) slice, 3 participants were missing the most proximal semimembranosus (SM) slice, and one participant was missing the two most proximal SM slices. Thus, these computations were made using fewer slices given that the muscle was not present at those anatomical levels on either leg at either timepoint.

### Muscle Biopsies

Mid-thigh VL biopsies were obtained using a 5-gauge needle with suction under aseptic conditions as previously described by our laboratory [13]. Approximately 80–140 mg of muscle tissue was excised per biopsy. Tissue allocated for histological analysis was mounted in optimal cutting temperature (OCT) compound, frozen in liquid nitrogen-cooled isopentane, and stored at −80°C. Tissue for molecular analyses (NEAT tissue) was rapidly cleaned of blood, fat, and connective tissue, placed in cryogenic vials, flash frozen in liquid nitrogen, and stored at −80°C. The tissue triage process took approximately 2 minutes for NEAT tissue and 4 minutes for OCT tissue.

### Immunohistochemistry

Cryosections (12 μm) of OCT tissue were cut using a cryostat (Leica Biosystems, Buffalo Grove, IL, USA) set to −22°C, adhered to positively charged histological slides, and stored at −80°C until batch processing. All samples from the same participant were processed on the same slide simultaneously.

On the day of immunohistochemistry, sections were retrieved from −80°C, air-dried for 2 hours at room temperature, and fixed in −20°C acetone for 5 minutes. Slides were then incubated with 3% H_2_O_2_ for 10 minutes, treated with autofluorescence quenching reagent (cat. no. 23007; Biotium, Fremont, CA, USA) for 1 minute and blocked for 1 hour with a solution of 5% goat serum, 2.5% horse serum, and 0.1% Triton-X at room temperature. Thereafter, slides were blocked with a streptavidin and biotin blocking solutions (cat. no. SP-2002; Vector Laboratories; Newark, CA, USA) at room temperature for 15 minutes each, with 5-minute phosphate-buffered saline (PBS) washes occurring between these incubations. Slides were then incubated overnight at 4°C with primary antibody cocktail containing 1:100 rabbit anti-dystrophin IgG (cat. no. GTX57970; Abcam; Waltham, MA, USA), 1:100 mouse anti-MyHCI IgG2b (clone: BA-D5; Developmental Studies Hybridoma Bank; Iowa City, IA, USA), and 1:20 mouse anti-PAX7 IgG1 (Developmental Studies Hybridoma Bank) + 2.5% horse serum in PBS. The following day, slides were washed four times (5 minutes each) in PBS and incubated for 90 minutes in biotin-SP-conjugated goat anti-mouse IgG1 (1:1,000) (cat. no. 111-065-003; Jackson ImmunoResearch; West Grove, PA, USA) in 2.5% BSA for 90 minutes. This step was followed by three 5-minute PBS washes and a 60-minute incubation with a solution containing 1:500 streptavidin (cat. no. S911; Thermo Fisher), 1:250 goat anti-mouse IgG2b Alexa Fluor 488 (cat. no. A21141; Thermo Fisher) and 1:250 goat anti-rabbit IgG Alexa Fluor 647 (cat. no. A21245; Thermo Fisher) in PBS. Following three 5-minute PBS washes, slides underwent a 20-minute incubation with 1:200 Alexa Fluor 555 tyramide (cat. no. B40955; Thermo Fisher) in PBS. Following three 5-minute PBS washes nuclei were stained with DAPI (1:10,000; cat. no. D3571; Thermo Fisher) for 10 minutes, washed twice (5 minutes each) in PBS, and sections were mounted with 1:1 PBS/glycerol. Digital images were acquired at 10× (fCSA) and 20× (myonuclei as well as satellite cells) objectives using a fluorescent microscope (Zeiss Axio Imager.M2 with motorized stage). Image analysis was automated using MyoVision v1.0 [14] for fiber type-specific fCSA and myonuclei per fiber. Satellite cells were manually counted by an investigator blinded to training conditions (C.A.E.) and expressed as number per 100 fibers (DAPI-positive & PAX7-positive cells within the dystrophin border).

### Total RNA Content

Approximately 20 mg of muscle tissue was homogenized in Trizol (cat. no. 15596026; Thermo Fisher) per manufacturer’s instructions. RNA pellets were reconstituted in 30 μL of diethyl pyrocarbonate (DEPC)-treated water and quantified in duplicate using a NanoDrop Lite spectrophotometer (Thermo Fisher). Total RNA content was normalized to input tissue wet weight and expressed as ng RNA per mg tissue, a validated surrogate of muscle ribosome content [15].

### Statistical Analysis

Aside from Experiment 1 transcriptomic analyses (described earlier in the Methods), all statistical analyses were conducted in R Studio using the lme4 and lmerTest packages. For Experiment 1 western blot outcomes and all Experiment 2 outcomes, a linear mixed-effects model (LMM) was fitted with Condition (FULL vs. LP), Time (PRE vs. POST for Experiment 2; PRE vs. 0h/3h/24h POST for Experiment 1), and their Condition × Time interaction as fixed effects. To account for the contralateral-limb design and repeated measurements, random intercepts were included for participant and for participant-by-condition, reflecting that each participant contributed both a FULL and LP limb measured at multiple points. Estimated marginal means, standard error (SE) values, 95% confidence intervals, and planned contrasts were derived using the emmeans package in R Studio [12]. For Experiment 1 western blot outcomes, follow-up pairwise comparisons between conditions at each timepoint were conducted with Sidak adjustment when a significant Condition × Time interaction was observed. For significant main effects of Time in the absence of a significant Condition × Time interaction, contrasts versus PRE collapsed across conditions were conducted with Sidak adjustment. Statistical significance was set at α = 0.05. However, interpretation focused on the magnitude and precision of estimated effects rather than statistical significance alone. Because equivalence margins were not prespecified, non-significant Condition × Time interactions were interpreted as indicating no clear evidence of a differential response between conditions, rather than formal statistical equivalence. To confirm the ROM manipulation, paired t-tests were used to compare FULL and LP conditions for knee and hip ROM and peak flexion angles at session 10, with each participant contributing one value per condition averaged across sets. Data are presented in-text as estimated marginal means with 95% confidence intervals (CI), and participant characteristics as well as data in figures are presented as mean ± standard deviation.

## RESULTS

### Participant Characteristics

Experiment 1 included 8 resistance-trained men (22 ± 1 years old, training experience of 5.6 ± 1 years) with a body mass of 92.4 ± 17.1 kg, DXA percent fat of 15.7 ± 5.6, and fat-free mass index of 21.0 ± 2.3 kg lean mass/m^2^, and no participant withdrew from the study. Experiment 2 enrolled 18 participants, of whom 2 withdrew (one before the intervention commenced for reasons unrelated to the study, and one due to knee pain), leaving 16 participants who completed the study. Participants were required to complete at least 14 of the 16 resistance training sessions, and none of the 16 finishers were below this threshold (average exercise adherence was 96.8%). The 16 finishers in Experiment 2 were 26 ± 5 years old (training experience of 8.0 ± 4.9 years) with a body mass of 86.3 ± 9.4 kg, BIS percent body fat of 18.2 ± 4.4, and fat-free mass index of 21.5 ± 1.2 kg lean mass/m^2^. For Experiment 2 biopsy outcomes, two participants were excluded from individual histological analyses given that tissue yield or quality was insufficient. Final sample sizes for fCSA, myonuclei, and SC analyses are reported in the results.

### Experiment 1: Acute Anabolic Phospho-signaling

Hippo and mTORC1 signaling protein data are presented in Figure 2. Phosphorylated LATS1 (Thr1079) demonstrated a significant Condition × Time interaction (P = 0.010), with post hoc analysis revealing FULL was greater than LP at 24h post (P = 0.004) (Fig. 2a). Phosphorylated YAP (Ser127) demonstrated a significant main effect of Time (P = 0.037), with values at 3h post lower than PRE (P = 0.015), but no Condition × Time interaction (P = 0.550) (Fig. 2b). Phosphorylated TAZ (Ser89) showed no significant main effect of Time (P = 0.249) and no Condition × Time interaction (P = 0.244) (Fig. 2c). For mTORC1 targets, phosphorylated mTOR (Ser2448) showed a significant main effect of Time (P = 0.010), with values at 0h post greater than PRE (P = 0.023), and no Condition × Time interaction (P = 0.787) (Fig. 2d). Phosphorylated p70S6K (Thr389) showed no significant main effect of Time (P = 0.231) and no Condition × Time interaction (P = 0.166) (Fig. 2e). Phosphorylated rpS6 (Ser235/236) demonstrated a significant main effect of Time (P = 0.033), with values at 0h post greater than PRE (P = 0.038), and no Condition × Time interaction (P = 0.353) (Fig. 2f). A representative Western blot for all targets is presented in Fig. 2g.

**FIGURE 2.**
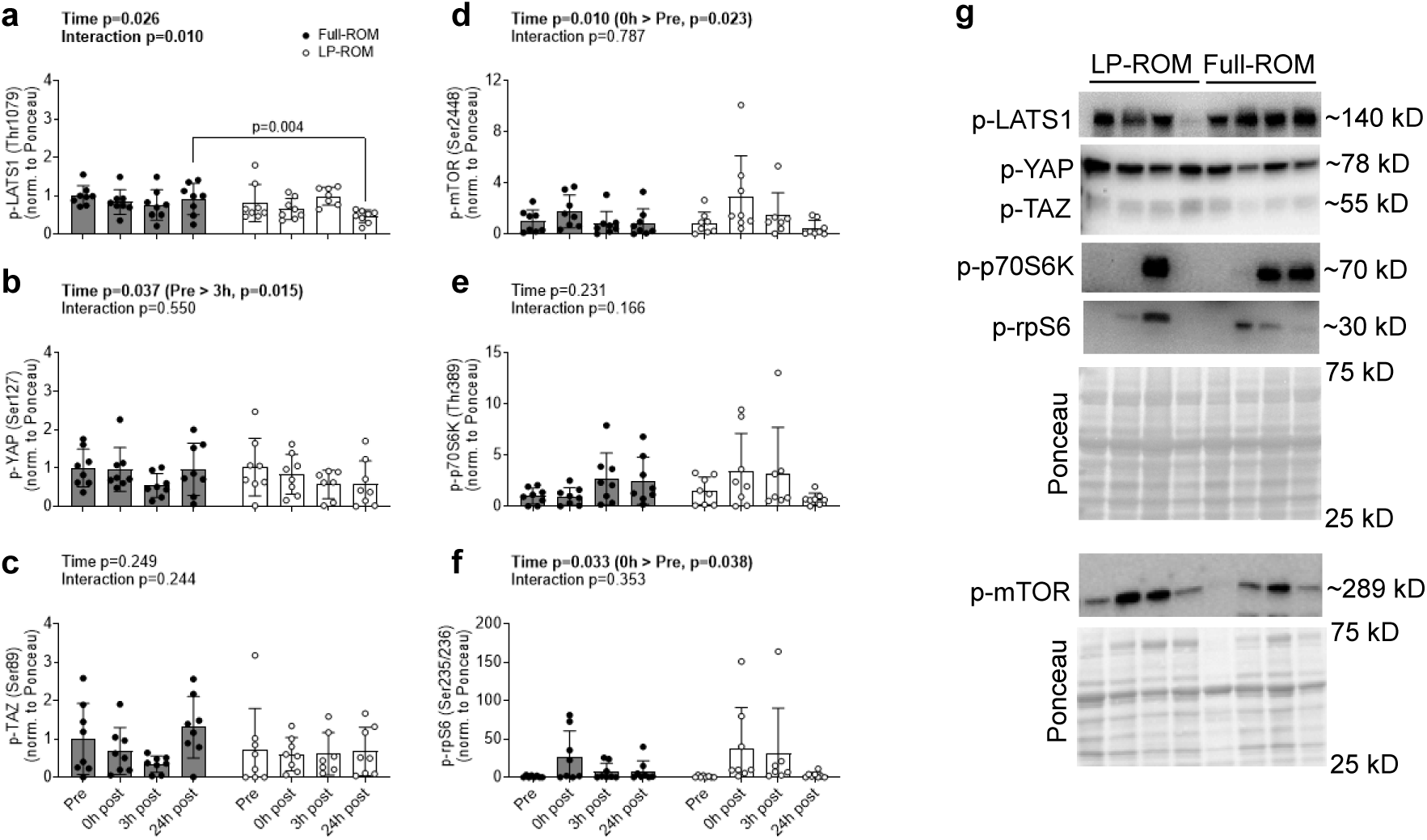
Experiment 1 Hippo and mTORC1 signaling protein data. Legend: Panels depict phosphorylated protein expression (normalized to Ponceau) for (a) p-LATS1 (Thr1079), (b) p-YAP (Ser127), (c) p-TAZ (Ser89), (d) p-mTOR (Ser2448), (e) p-p70S6K (Thr389), and (f) p-rpS6 (Ser235/236) in the VL of n=8 participants at PRE, 0h, 3h, and 24h post-exercise following Full-ROM (filled circles) and LP-ROM (open circles) leg extension bouts. Representative western blot images and Ponceau stain are also presented. Time and Condition × Time interaction P values from LMMs are shown above each panel; bolded interaction P values indicate P < 0.05. Bars represent means ± standard deviation values with individual participant values are shown as dots.

### Experiment 1: Acute Transcriptomic Responses

Both LP and FULL conditions produced progressive increases in the number of differentially expressed mRNAs relative to PRE across the post-exercise time course. Upregulated mRNAs relative to PRE were 31 (FULL) and 18 (LP) at 0h, 707 and 469 at 3h, and 1,344 and 1,242 at 24h post-exercise (Fig. 3a). Downregulated mRNAs were 9 (FULL) and 8 (LP) at 0h, 157 and 76 at 3h, and 1,379 and 975 at 24h post-exercise (Fig. 3b). mRNAs showing significant interactions were 22 (up in FULL) and 9 (up in LP) from Pre to 0h, 20 (up in FULL) and 24 (up in LP) from Pre to 3h, and were 35 (up in FULL) and 5 (up in LP) from Pre to 24h post-exercise (Fig. 3c). Enriched Gene Ontology (GO) pathways from upregulated mRNAs within-condition were absent from Pre to 0h in both conditions, with 34 (up in FULL) and 15 (up in LP) enriched pathways from Pre to 3h, and 38 (up in FULL) and 50 (up in LP) Pre to 24h post-exercise (Fig. 3d). Minimal GO pathway enrichment was observed from downregulated mRNAs (Fig. 3e). Moreover, no enriched pathways were evident from the limited number of mRNAs (seen in Fig. 3c) exhibiting significant interactions (Fig. 3f).

**FIGURE 3.**
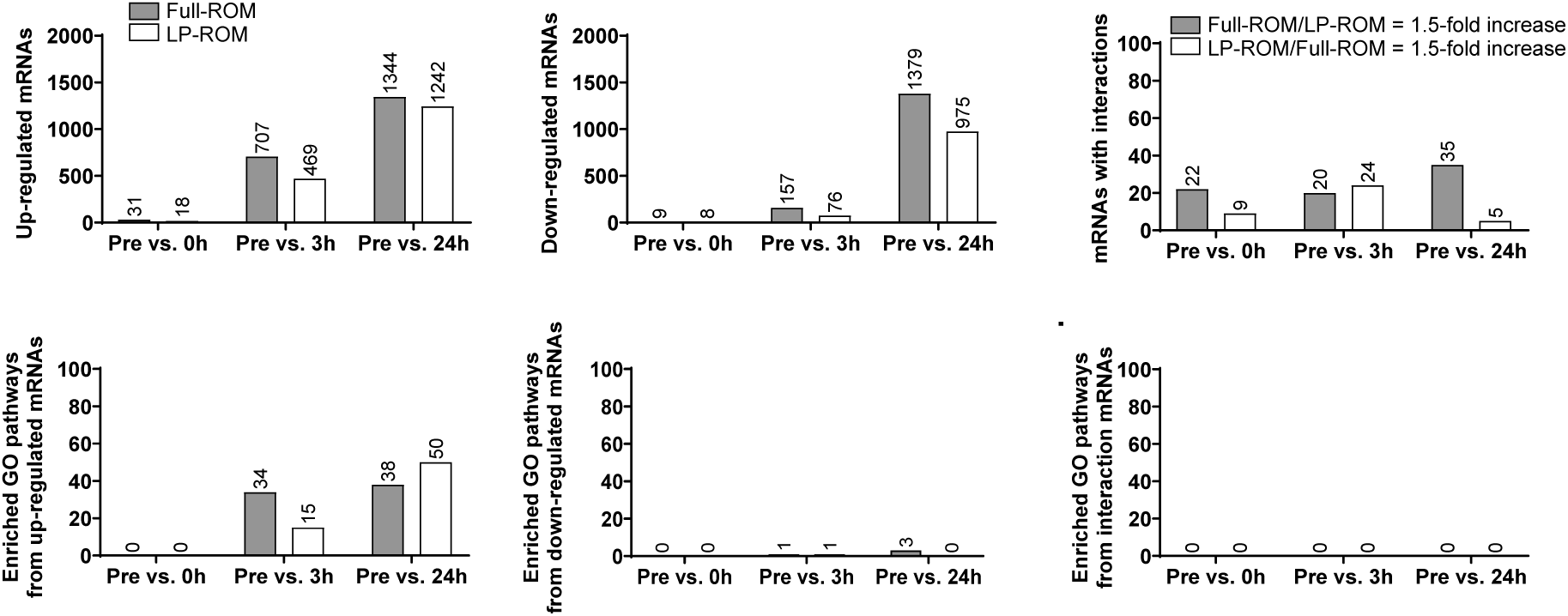
Experiment 1 summary of transcriptomic data. Legend: Full-ROM (gray bars) and LP-ROM (open bars) leg extension bouts in the VL of n=8 resistance-trained men. Panels depict (a) the number of up-regulated mRNAs within each condition relative to Pre, (b) down-regulated mRNAs within each condition relative to Pre, (c) mRNAs exhibiting between-condition interactions, (d) enriched Gene Ontology (GO)-Slim biological pathways from up-regulated mRNAs within each condition, (e) enriched GO-Slim pathways from down-regulated mRNAs within each condition, and (f) enriched GO-Slim pathways from mRNAs demonstrating interactions.

Figure 4 provides additional context related to the within-leg protocol transcriptomic changes from Pre to 3h post-exercise (Fig. 4a&b) as well as Pre to 24h post-exercise (Fig. 4c&d). Two conserved GO-Slim biological pathways were evident within protocols based on upregulated mRNAs from Pre to 3h post-exercise and included “inflammatory response” (GO:0006954) and “regulation of angiogenesis” (GO:0045765). Moreover, the transcriptional response was largely conserved between conditions, with 13 of the top 20 upregulated mRNAs from PRE to 3h post-exercise and 14 of the top 20 upregulated mRNAs from PRE to 24h post-exercise shared across LP and FULL protocols (ranked by fold-change).

**FIGURE 4.**
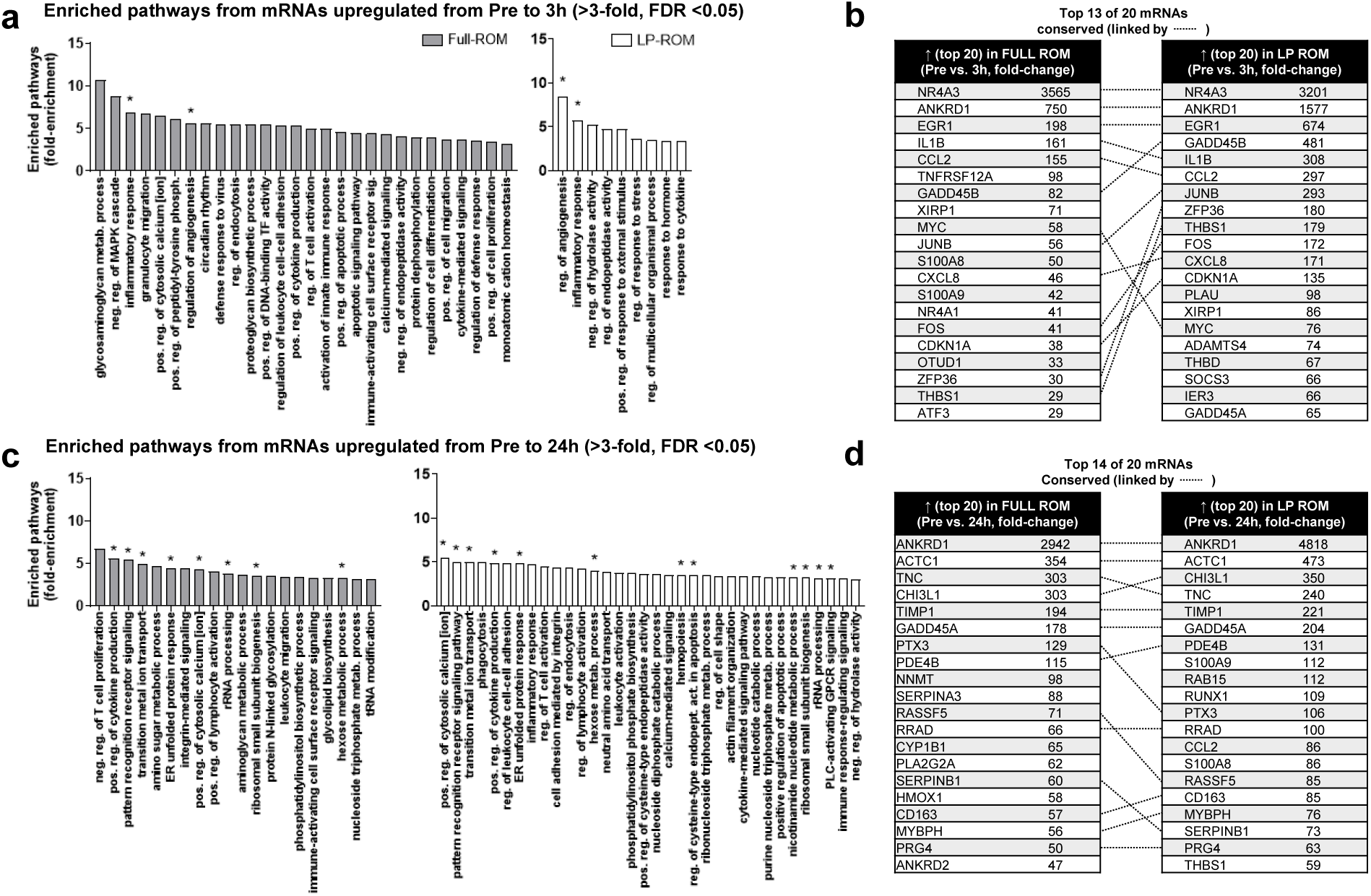
Experiment 1 details of enriched GO-Slim biological pathways and top up-regulated mRNAs within each condition. Legend: Full-ROM (gray bars) and LP-ROM (open bars) leg extension bouts in the VL of n=8 resistance-trained men. Panels depict (a) the top GO-Slim biological pathways predicted to be enriched based on up-regulated mRNAs in both protocols from Pre to 3h post-exercise, (b) the top 20 up-regulated mRNAs in both protocols from Pre to 3h post-exercise, (c) the top GO-Slim biological pathways predicted to be enriched based on up-regulated mRNAs in both protocols from Pre to 24h post-exercise, (d) up-regulated mRNAs in both protocols from Pre to 24h post-exercise. Note: in panels a & c only >3-fold enriched pathways (FDR < 0.05) are presented due to graphical size restrictions. Symbols: * in panels a & c depict enriched pathways (FDR < 0.05) that were conserved between protocols at the Pre to 3h or Pre to 24h comparisons, and dashed lines (--------) depict conserved mRNAs between protocols in panels b&d.

### Experiment 2: Kinematics

Peak and minimum joint angles during the leg press exercise, measured at session 10, are presented in Figure 5, with individual participant data shown for each condition. For the hip, peak flexion was similar between conditions (FULL: 103.9 ± 13.2°; LP: 106.8 ± 9.2°; P = 0.094), while minimum hip flexion differed markedly (FULL: 62.1 ± 12.1°; LP: 85.8 ± 9.8°), resulting in a substantially greater hip ROM in FULL compared to LP (41.8 ± 7.7° vs. 21.0 ± 3.4°; t(14) = 10.45, P < 0.001). For the knee, peak flexion was again similar between conditions (FULL: 123.7 ± 11.7°; LP: 126.2 ± 12.6°; P = 0.328), while minimum knee flexion differed markedly (FULL: 31.0 ± 10.4°; LP: 78.3 ± 10.8°), resulting in a substantially greater knee ROM in FULL compared to LP (92.7 ± 10.5° vs. 47.9 ± 8.7°; t(14) = 15.44, P < 0.001). Collectively, these data confirm that the LP condition achieved approximately half the ROM of FULL at both the knee and hip, consistent with the intended manipulation.

**FIGURE 5.**
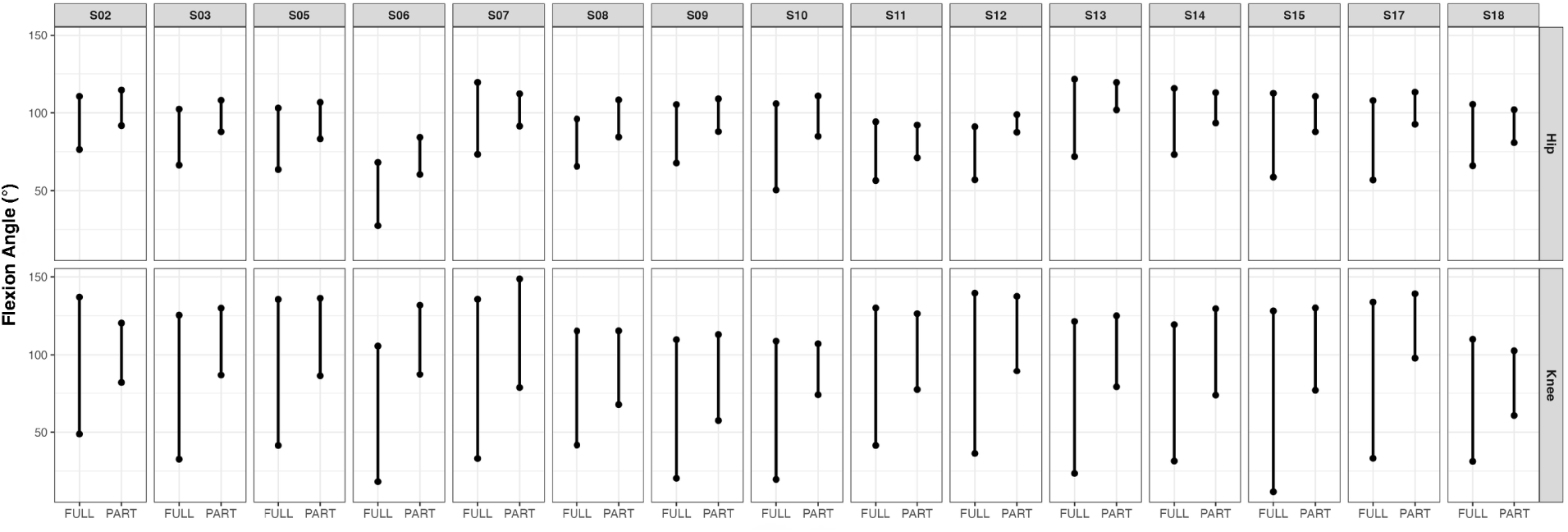
Experiment 2 knee and hip flexion angle ranges during the leg press exercise recorded at session 10. Legend: Each panel displays individual participant knee and hip flexion kinematics data (n=15) recorded for a single Full-ROM (FULL) and lengthened partial-ROM (LP) bout (session 10 of 16). Filled circles represent peak (upper) and minimum (lower) flexion angles averaged across sets within the session.

### Experiment 2: VL Muscle Cross-Sectional Area

VL mCSA data are presented in Figure 6. Total VL mCSA, summed across five equidistant MRI-derived transverse slices increased from PRE to POST when conditions were collapsed, indicating evidence of VL hypertrophy over the intervention period (main effect of Time estimate: 9.3 cm^2^, 95% CI [6.8, 11.8], P < 0.001), FULL increased from 166.0 ± 6.7 to 175.9 ± 6.7 cm^2^, and LP increased from 164.7 ± 6.7 to 173.4 ± 6.7 cm^2^. There was no clear evidence of a differential change between conditions (Condition × Time contrast estimate: −1.4 cm², 95% CI [−6.08, 3.8], P = 0.640).

**FIGURE 6.**
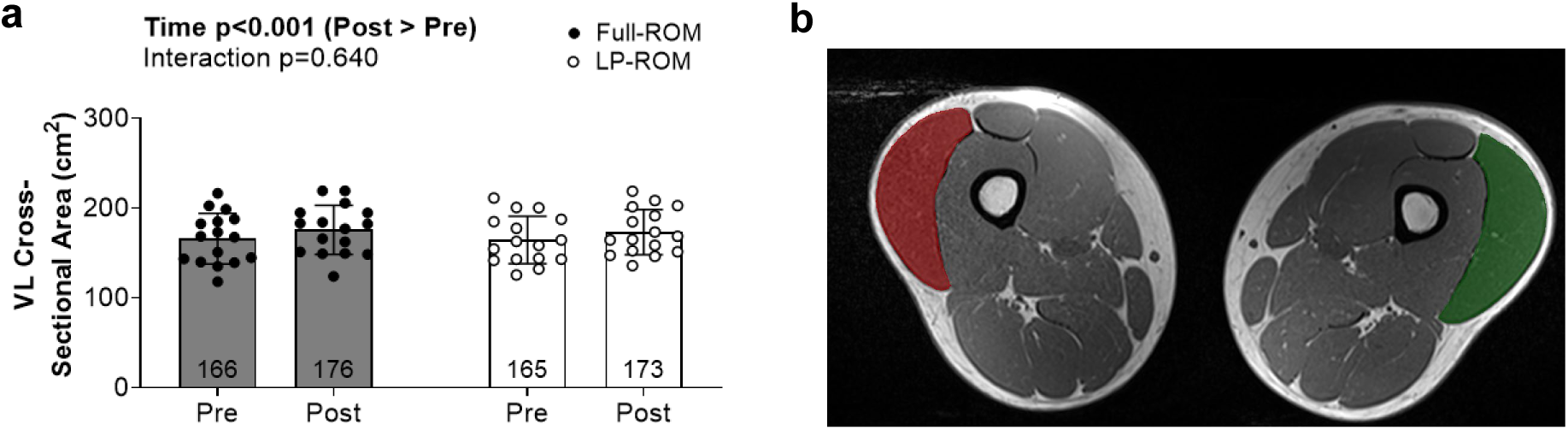
Experiment 2 VL muscle cross-sectional area data. Legend: Results are from n=16 participants. Panels depict (a) total VL mCSA summed across five equidistant MRI-derived transverse slices at PRE and POST for Full-ROM (filled circles) and LP-ROM (open circles) training conditions. Bars represent means ± standard deviation values with individual participant values are shown as dots. Time and Condition × Time interaction P values from LMMs are shown above. (b) Representative transverse MRI images showing VL segmentation (red = FULL leg PRE; green = LP leg POST) at mid-thigh from the same participant.

### Experiment 2: Fiber Type-Specific Cross-Sectional Area, and Myonuclei

Type I fCSA and myonuclei per fiber data are presented in Fig. 7a&c. Type I fCSA did not provide clear evidence of an overall PRE-to-POST change when conditions were collapsed (main effect of Time estimate: 241 μm², 95% CI [−242, 724], P = 0.314). Estimated marginal means were 5,123 ± 318 μm² at PRE and 5,194 ± 318 μm² at POST in FULL, and 5,280 ± 318 μm² at PRE and 5,703 ± 327 μm² at POST in LP. There was no clear evidence of a differential change between conditions (Condition × Time contrast estimate: 339 μm², 95% CI [−627, 1305], P = 0.476). Similarly, Type I myonuclei per fiber did not provide clear evidence of an overall PRE-to-POST change when conditions were collapsed (main effect of Time estimate: −0.22, 95% CI [−0.60, 0.16], P = 0.245). Estimated marginal means were 2.39 ± 0.19 at PRE and 2.13 ± 0.19 at POST in FULL, and 2.35 ± 0.20 at PRE and 2.18 ± 0.20 at POST in LP. There was no clear evidence of a differential change between conditions (Condition × Time contrast estimate: 0.09, 95% CI [−0.67, 0.85], P = 0.813).

**FIGURE 7.**
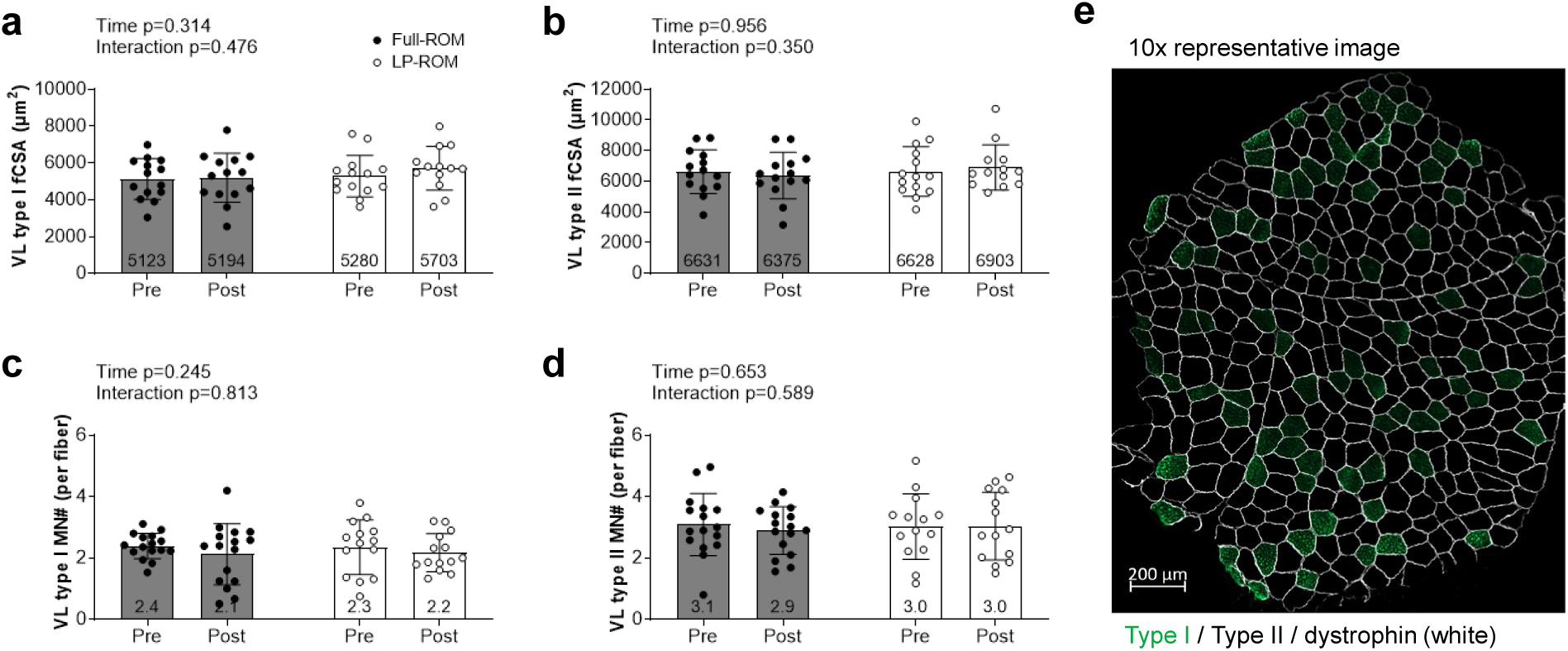
Experiment 2 VL fiber cross-sectional area and myonuclei data. Legend: Panels depict (a) Type I fCSA, (b) type II fCSA, (c) type I myonuclei per fiber, and (d) type II myonuclei per fiber at PRE and POST for Full-ROM (filled circles) and LP-ROM (open circles) conditions. Data are from 14 of the 16 participants due to two participants not yielding tissue quantity or quality sufficient for analyses. Bars represent means ± standard deviation values with individual participant values are shown as dots. Time and Condition × Time interaction P values are shown above each panel. (e) Representative 10× fluorescence microscopy image from one participant showing type I fibers (Alexa Fluor 488), type II fibers (not stained), and dystrophin (Alexa Fluor 647, pseudo-colored white border).

Type II fCSA and myonuclei per fiber data are presented in Fig. 7b&d. Type II fCSA did not provide clear evidence of an overall PRE-to-POST change when conditions were collapsed (main effect of Time estimate: 14.5 μm², 95% CI [−569, 599], P = 0.956). Estimated marginal means were 6,630 ± 402 μm² at PRE and 6,374 ± 402 μm² at POST in FULL, and 6,628 ± 401 μm² at PRE and 6,913 ± 413 μm² at POST in LP. There was no clear evidence of a differential change between conditions (Condition × Time contrast estimate: 541 μm², 95% CI [−627, 1709], *P* = 0.350). Similarly, Type II myonuclei per fiber did not provide clear evidence of an overall PRE-to-POST change when conditions were collapsed (main effect of Time estimate: −0.09, 95% CI [−0.49, 0.31], P = 0.653). Estimated marginal means were 3.09 ± 0.25 at PRE and 2.89 ± 0.25 at POST in FULL, and 3.02 ± 0.26 at PRE and 3.03 ± 0.26 at POST in LP. There was no clear evidence of a differential change between conditions (Condition × Time contrast estimate: 0.21, 95% CI [−0.59, 1.01], P = 0.589).

### Experiment 2: Satellite Cell Number, and Total RNA Content

Satellite cell and total RNA data are presented in Figure 8. Type I satellite cell number did not provide clear evidence of an overall PRE-to-POST change when conditions were collapsed (main effect of Time estimate: 1.9, 95% CI [−1.0, 4.7], P = 0.186). Estimated marginal means were 9.1 ± 3.0 at PRE and 8.7 ± 3.0 at POST in FULL, and 8.8 ± 3.0 at PRE and 13.1 ± 3.1 at POST in LP. There was no clear evidence of a differential change between conditions (Condition × Time contrast estimate: 4.7, 95% CI [−1.0, 10.3], P = 0.102). In contrast, Type II satellite cell number increased from PRE to POST when conditions were collapsed, indicating evidence of satellite cell accretion over the intervention period (main effect of Time estimate: 8.3, 95% CI [2.2, 14.4], P = 0.001); however, there was no clear evidence of a differential change between conditions (Condition × Time contrast estimate: 1.6, 95% CI [−10.7, 13.8], P = 0.797), and estimated marginal means were 11.5 ± 3.9 at PRE and 19.0 ± 3.9 at POST in FULL, and 7.8 ± 3.9 at PRE and 16.9 ± 4.0 at POST in LP.

**FIGURE 8.**
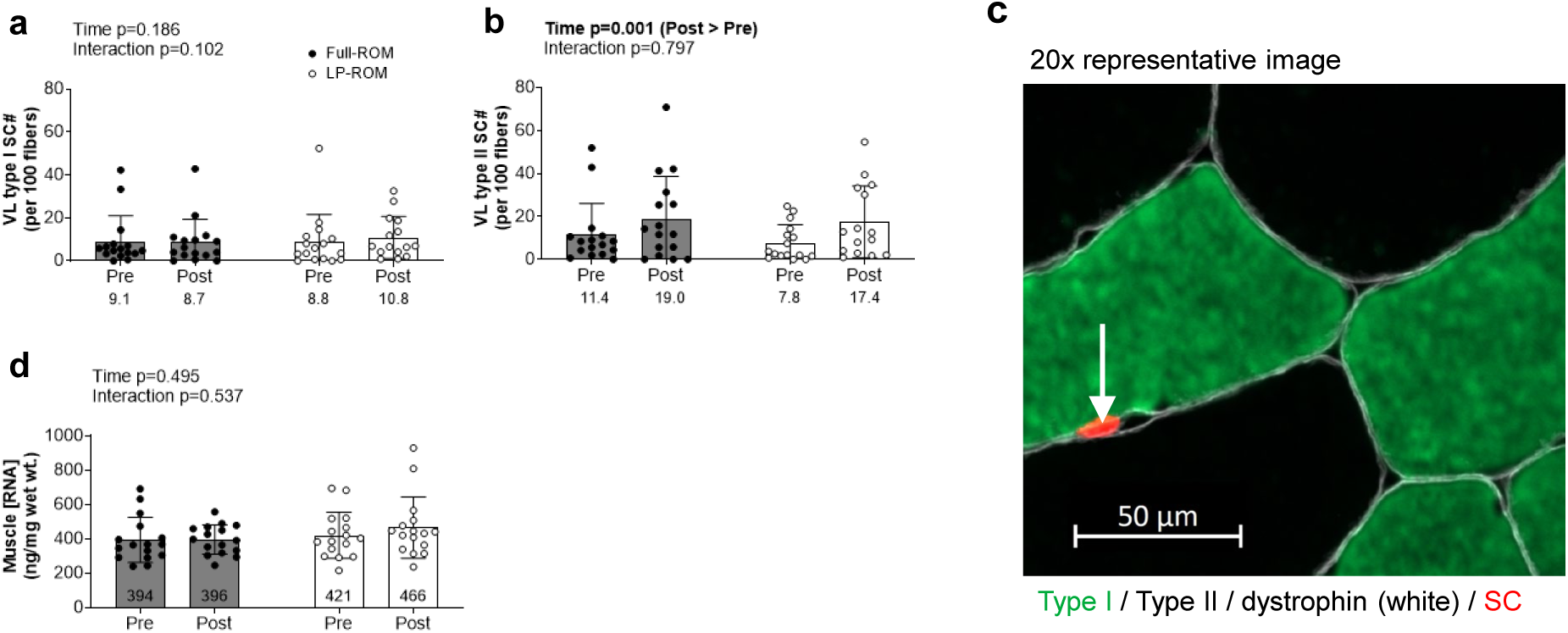
Experiment 2 VL satellite cell number and total RNA content. Legend: Panels depict (a) Type I SC per 100 fibers and (b) type II SC per 100 fibers at PRE and POST for Full-ROM (filled circles) and LP-ROM (open circles) conditions. Data are from 14 participants for satellite cells and 16 for total RNA. (c) Representative 20× fluorescence microscopy image from one participant showing type I/type II fibers, dystrophin (pseudocolored white border), and a type I fiber satellite cell (within the dystrophin border). (d) Total RNA content (ng/mg wet tissue) at PRE and POST. Bars represent means ± standard deviation values with individual participant values are shown as dots. Time and Condition × Time interaction P values are shown above each panel.

Total RNA content did not provide clear evidence of an overall PRE-to-POST change when conditions were collapsed (main effect of Time estimate: 23.5 ng/mg, 95% CI [−45.9, 93.0], P = 0.495). Estimated marginal means were 394 ± 34 ng/mg at PRE and 396 ± 34 ng/mg at POST in FULL, and 421 ± 34 ng/mg at PRE and 465 ± 34 ng/mg at POST in LP. There was no clear evidence of a differential change between conditions (Condition × Time contrast estimate: 42.5 ng/mg, 95% CI [−96.4, 181.0], P = 0.537).

### Experiment 2: Cross-Sectional Area Values of Non-VL Muscles

Experiment 2 mCSA data for all thigh and hip muscles are presented in Figure 9 and Table 1. The PRE-to-POST changes in mCSA values for the adductors, the remaining three quadriceps muscles (rectus femoris, vastus intermedius, vastus medialis), gracilis, sartorius, biceps femoris long head, and semimembranosus did not show clear evidence favoring one protocol. In contrast, semitendinosus mCSA showed greater estimated change following LP than FULL training (Condition × Time contrast estimate: 2.4 cm², 95% CI [0.5, 4.3], P = 0.015), while summed hamstring mCSA changes showed a similar directional pattern, although the 95% CI included zero (Condition × Time contrast estimate: 3.9 cm², 95% CI [−0.2, 7.9], P = 0.058).

**FIGURE 9.**
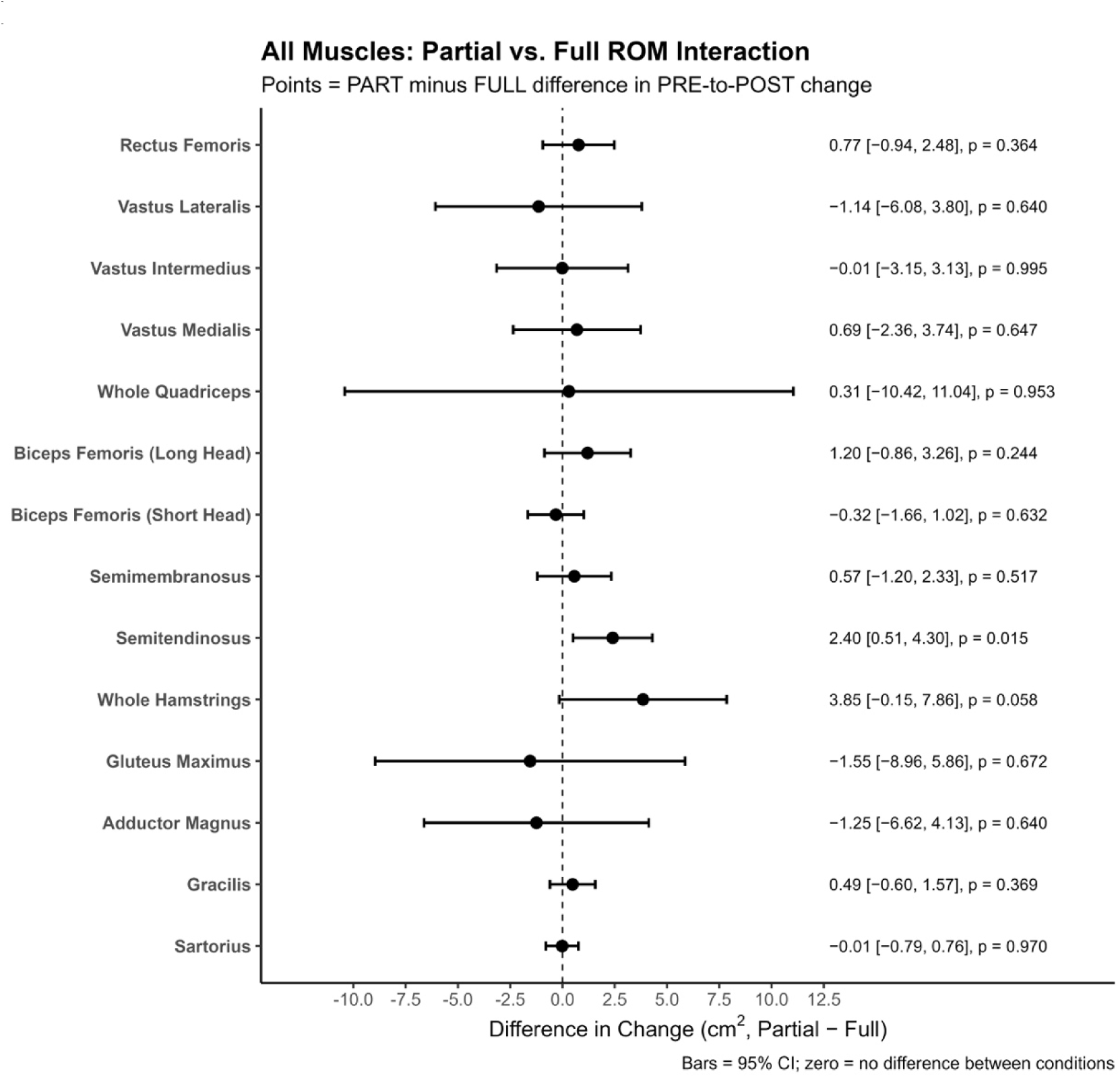
Experiment 2 secondary MRI analysis of all muscles. Legend: Results are from n=16 participants for all muscles except gluteus maximus (n=15). Condition × Time contrast estimates and 95% CIs are shown for the rectus femoris, vastus lateralis, vastus intermedius, vastus medialis, whole quadriceps, biceps femoris long head, biceps femoris short head, semimembranosus, semitendinosus, whole hamstrings, gluteus maximus, adductors, gracilis, and sartorius. Positive values indicate greater change in the LP condition relative to FULL. The dashed vertical line represents zero (no difference between conditions).

**TABLE 1.** Experiment 2 secondary MRI analysis muscle cross-sectional area values by condition.

| Muscle | Full-ROM |  |  | LP-ROM |  |  |
| --- | --- | --- | --- | --- | --- | --- |
| | *Pre* | *Post* | $\Delta$ | *Pre* | *Post* | $\Delta$ |
| ***Quadriceps*** |  |  |  |  |  |  |
| RF | 39.3 ± 7.3 | 41.1 ± 7.6 | +1.8 ± 2.0 | 38.6 ± 7.7 | 41.2 ± 8.5 | +2.6 ± 2.7 |
| VI | 138.0 ± 18.1 | 144.3 ± 18.6 | +6.2 ± 4.7 | 135.1 ± 18.8 | 141.3 ± 18.7 | +6.2 ± 4.0 |
| VL | 166.0 ± 28.3 | 175.9 ± 27.4 | +9.9 ± 6.7 | 164.7 ± 26.4 | 173.4 ± 25.2 | +8.7 ± 7.0 |
| VM | 120.1 ± 16.4 | 125.5 ± 14.3 | +5.4 ± 4.3 | 117.5 ± 18.5 | 123.6 ± 17.1 | +6.1 ± 4.1 |
| WQ | 463.4 ± 57.0 | 486.8 ± 52.6 | +23.4 ± 14.7 | 455.9 ± 58.7 | 479.5 ± 54.8 | +23.7 ± 15.0 |
| ***Hamstrings*** |  |  |  |  |  |  |
| BFlh | 68.1 ± 8.5 | 70.0 ± 8.0 | +1.9 ± 3.2 | 66.0 ± 8.9 | 69.0 ± 8.7 | +3.1 ± 2.5 |
| BFsh | 21.2 ± 5.8 | 23.3 ± 5.9 | +2.1 ± 1.9 | 21.7 ± 6.4 | 23.5 ± 5.8 | +1.8 ± 1.8 |
| SM | 50.5 ± 9.2 | 50.8 ± 9.2 | +0.3 ± 2.2 | 49.7 ± 10.4 | 50.6 ± 9.5 | +0.9 ± 2.7 |
| ST | 49.8 ± 10.9 | 52.6 ± 10.9 | +2.8 ± 2.3 | 51.6 ± 10.4 | 56.8 ± 10.6 | +5.2 ± 2.9 |
| WH | 189.6 ± 21.3 | 196.7 ± 18.5 | +7.1 ± 5.5 | 189.0 ± 26.2 | 199.9 ± 22.7 | +10.9 ± 5.6 |
| ***Other*** |  |  |  |  |  |  |
| SA | 24.7 ± 4.1 | 26.0 ± 4.2 | +1.3 ± 0.7 | 24.6 ± 4.4 | 25.8 ± 4.4 | +1.3 ± 1.3 |
| GR | 26.0 ± 5.5 | 27.5 ± 6.1 | +1.5 ± 1.5 | 26.1 ± 4.8 | 28.1 ± 4.9 | +2.0 ± 1.5 |
| ADD | 172.6 ± 37.0 | 180.5 ± 40.1 | +7.9 ± 8.2 | 172.8 ± 32.0 | 179.4 ± 33.4 | +6.7 ± 6.6 |
| GM | 217.4 ± 21.5 | 231.1 ± 22.1 | +13.7 ± 8.6 | 212.9 ± 18.6 | 225.1 ± 19.2 | +12.1 ± 11.0 |

Legend: Values are mean ± standard deviation (cm<sup>2</sup>). Symbol: $\Delta$ , Post minus Pre values. Abbreviations : RF, rectus femoris; VI, vastus intermedius; VL, vastus lateralis; VM, vastus medialis; WQ, whole quadriceps; BFlh, biceps femoris long head; BFsh, biceps femoris short head; SM, semimembranosus; ST, semitendinosus; WH, whole hamstrings; SA, sartorius; GR, gracilis; ADD, adductors; GM, gluteus maximus.

## DISCUSSION

The primary finding of this investigation is that 8 weeks of LP and FULL lower-body resistance training produced increases in VL size, with no clear evidence that hypertrophic or cellular responses differed between conditions in resistance-trained men. Summed VL mCSA values across five MRI-derived equidistant images increased significantly with training in both conditions, confirming that both protocols elicited hypertrophy. However, Condition × Time contrasts did not provide clear evidence of differential change between LP and FULL for any primary outcome, including VL mCSA, fiber type-specific CSA, myonuclei per fiber, satellite cell number, or total RNA content. In the accompanying acute study, both conditions produced broadly similar time-dependent VL transcriptomic changes and anabolic signaling responses. However, secondary MRI analyses provided some evidence of a possible LP advantage for hamstring hypertrophy, particularly for the semitendinosus, with summed hamstring mCSA showing a similar but less certain directional pattern. The following paragraphs expand upon these findings.

In Experiment 1, both LP and FULL protocols produced remarkably conserved acute transcriptional and signaling responses in the VL. This was evident through similar enriched GO-Slim biological pathways from upregulated mRNAs shared across conditions at 3h and 24h post-exercise (as well as conserved top-upregulated mRNAs themselves). Regardless of protocol, these pathway-level findings are broadly consistent with the resistance exercise transcriptomics literature. The MetaMEx meta-analysis conducted by Pillon and colleagues integrating data from 66 skeletal muscle transcriptomic datasets, including 8 studies of acute resistance exercise, identified the inflammatory and angiogenesis responses as one of the most consistently enriched GO terms among upregulated genes in the 2–4h window following acute resistance exercise in healthy humans [16]. A subsequent analysis from the same group employing the MetaMEx database to construct a transcriptional timeline from 0 to 48h after acute exercise revealed that inflammatory response and cytokine signaling pathways are characteristically induced in skeletal muscle [17], mirroring the 24h timepoint at which these pathways emerged in the present study. Importantly, the presence of the inflammatory response in these resistance-trained participants should not be interpreted solely as a marker of exercise-induced muscle damage. Rather, the transcriptional upregulation of inflammatory mediators likely reflects regulated remodeling that facilitates satellite cell recruitment, extracellular matrix turnover, and protein turnover, and these responses notably persist in experienced trainees as sampled in the current study [16]. The Hippo and mTORC1 signaling data also largely showed no acute differences between-condition over time. The one significant Condition × Time interaction observed was for phosphorylated LATS1 (Thr1079), a component of the Hippo signaling pathway, which was higher in FULL relative to LP at the 24-hour post-exercise timepoint (P = 0.004). The Hippo pathway has been implicated in mechanosensory transduction and load-dependent gene regulation and LATS1 phosphorylation at Thr1079 reflects kinase activation that generally suppresses downstream YAP/TAZ activity [18]. However, the biological significance of this isolated Experiment 1 interaction is difficult to interpret given the lack of any other Experiment 2 (chronic) interactions. Hence, the Experiment 1 collective outcomes suggest both conditions likely impose a qualitatively comparable acute mechanical challenge on the VL muscle.

In Experiment 2, the absence of significant training effects on fiber type-specific fCSA in either condition also warrants consideration. Overall VL mCSA increased significantly, though modestly, confirming that whole-muscle hypertrophy occurred. The MRI finding is likely related to resistance training in previously trained individuals producing smaller relative increases in skeletal muscle hypertrophy compared to untrained populations [19]. Further, 8 weeks at two sessions per week may be insufficient to detect significant fiber-level hypertrophy due to the small tissue sampling with biopsy analysis along with variation in this measure as our laboratory and others have previously posited in prior reviews and original investigations [20–22]. It is also worth noting that the lack of biopsy-derived fCSA increases in the context of VL hypertrophy likely reflects a localized sample at a fixed mid-thigh site that may not capture the regional distribution of hypertrophy across the muscle as comprehensively as MRI. The multi-site VL MRI data, therefore, provide a more sensitive and anatomically complete estimate of hypertrophic response in this study.

Both Experiment 2 protocols also produced a significant increase in type II satellite cell number over the intervention, with no clear evidence that this response differed between LP and FULL. Satellite cells are essential for long-term muscle fiber remodeling and their expansion with resistance training is well-established in trained populations [23, 24]. The similar pattern of satellite cell responses observed between LP and FULL further supports the conclusion that ROM does not fundamentally alter the cellular remodeling environment in the VL. Total RNA content, a validated surrogate of muscle ribosome content, did not increase significantly in either condition. This finding is consistent with prior data from our laboratory indicating that shorter-term resistance training programs in individuals with prior training experience do not increase this outcome [25]. Moreover, the substantial variability in this measure, evidenced by wide confidence intervals, may also have limited the ability to detect a training effect. Regardless, the absence of a condition difference in total RNA reinforces that LP and FULL imposed a comparable stimulus with respect to translational machinery remodeling in the VL.

Critically, a biomechanical rationale may help explain the null VL findings across both experiments. During Experiment 2, deep knee flexion in the FULL ROM leg press positions the VL in a more lengthened state while substantial external moment demand is present at the joint [26], meaning conventional FULL training already imposes considerable mechanical challenge at longer muscle lengths. Although a full ROM leg extension performed on a conventional weight stack machine presents a shortened-dominant resistance challenge to the VL, its inclusion alongside the leg press within a multi-exercise program may have rendered it a negligible contributor to the overall hypertrophic stimulus. Findings from both experiments are broadly consistent with prior work in the quadriceps suggesting that VL hypertrophy is not strongly influenced by ROM differences within a practically relevant range. For instance, interrogations into squat depth indicate that knee flexion beyond approximately 90° does not further enhance VL hypertrophy compared to shallower depths [27], and the present Experiment 2 data extend this to a direct LP versus FULL comparison across multiple lower-body exercises in trained individuals.

Contrary to the largely null VL data, our Experiment 2 secondary MRI analyses did indicate differences exist between training protocols that warrant further discussion. To our knowledge, this represents the first study to examine the effects of partial versus full ROM training on hamstring hypertrophy directly. Prior work has demonstrated that biarticular hamstring hypertrophy is sensitive to muscle length, with greater growth observed when trained in hip-flexed versus hip-neutral positions [4, 5], suggesting a length-dependent advantage. However, those studies manipulated joint position rather than ROM. The present findings extend this by suggesting that LP training, which emphasizes the lengthened portion of the movement arc, may confer a hamstring hypertrophy advantage over FULL training. It is worth noting that the lying leg curl was the only exercise targeting the hamstrings directly, and the lying leg curl utilized in the present study presents a shortened-dominant resistance challenge. Training this exercise through the lengthened portion of the ROM therefore shifted the mechanical challenge toward longer muscle lengths, which could explain the observed LP advantage. Among individual hamstring muscles, the semitendinosus demonstrated the greatest absolute hypertrophy, which may have provided sufficient statistical power to detect a between-condition difference that was present but undetected in muscles exhibiting more modest overall growth, with the summed hamstring analysis also providing directional support for LP. Collectively, these findings suggest that the advantage of lengthened-biased training may be most apparent when a muscle is trained with more limited exercise selection against a shortened-dominant resistance challenge, as the contrast between LP and FULL conditions is greatest under these circumstances. However, whether this pattern generalizes across muscles remains unclear and warrants further investigation.

### Experimental considerations

Several limitations should be acknowledged. The high costs of -omics analyses (Experiment 1) and MRI (Experiment 2) precluded the inclusion of a large participant pool in either study. Notwithstanding, this was somewhat mitigated using within-subject designs for both studies, which reduces between-participant variability [28]. Second, we did not measure sarcomere length, serial sarcomere number, or fascicle length adaptations in Experiment 2, which would have provided additional mechanistic insight into potential architectural adaptations to both protocols. Third, only younger trained men were examined herein, and these results do not reflect potential adaptations that may be observed in other populations.

## Conclusions

Despite the aforementioned limitations, this is the first study to extensively examine the acute-molecular and chronic cellular adaptations to LP resistance training. Both LP and FULL lower-body protocols resulted in increased VL hypertrophy and similar patterns of cellular adaptations over 8 weeks in resistance-trained men, as assessed by MRI-derived CSA, fiber type-specific CSA, myonuclei per fiber, SC number, and total RNA content. Experiment 1 acute transcriptomic and anabolic signaling responses were similarly comparable between LP and FULL protocols, which lend further credence to the chronic VL outcomes observed in Experiment 2. Notably, secondary analyses suggested that LP training may confer a hamstring hypertrophy advantage over FULL training, potentially reflecting the shortened-dominant resistance challenge of the lying leg curl exercise employed in the present study. Future work should attempt to replicate findings and continue to characterize the muscle-specific boundary conditions and contexts under which length-dependent hypertrophy is more robustly observed, which may have practical implications for those who are time or exercise selection constrained. Importantly, the present findings suggest that within the context of a multi-exercise resistance training program incorporating exercises that challenge muscles across varied portions of the ROM, the advantage of lengthened-biased training may be attenuated or absent.

## ADDITIONAL INFORMATION

## Acknowledgments

We graciously thank the participants for partaking in this study.

## Research ethics

Experiment 1 was approved by the University of Mary Hardin-Baylor’s IRB (IRB protocol no. LT81725). Experiment 2 was approved by the Auburn University IRB (IRB protocol no. STUDY00000282).

## Informed consent statement

All participants in Experiments 1 and 2 were provided with ample information about study procedures and provided verbal and written consent prior to engaging in protocols.

## Authorship contribution statement

DLP primarily drafted the manuscript with the assistance of MDR. DRT, TG, JK, MR, JQ, CDW, MM-V, VR-C, AB, RJB, CAE, and CGV aided in multiple aspects of data collection and/or analysis warranting co-authorship. LWT supervised and oversaw Experiment 1 execution. This project served as DLP’s dissertation works, and project supervision was provided by CBM, RB, ANK, and DTB. All co-authors were provided with the opportunity to edit the work and agree to the manuscript’s submission.

## Data availability

Raw data related to the study outcomes will be provided upon reasonable request by emailing.

